# Random versus maximum entropy models of neural population activity

**DOI:** 10.1101/092973

**Authors:** Ulisse Ferrari, Tomoyuki Obuchi, Thierry Mora

## Abstract

The principle of maximum entropy provides a useful method for inferring statistical mechanics models from observations in correlated systems, and is widely used in a variety of fields where accurate data are available. While the assumptions underlying maximum entropy are intuitive and appealing, its adequacy for describing complex empirical data has been little studied in comparison to alternative approaches. Here data from the collective spiking activity of retinal neurons is reanalysed. The accuracy of the maximum entropy distribution constrained by mean firing rates and pairwise correlations is compared to a random ensemble of distributions constrained by the same observables. In general, maximum entropy approximates the true distribution better than the typical or mean distribution from that ensemble. This advantage improves with population size, with groups as small as 8 being almost always better described by maximum entropy. Failure of maximum entropy to outperform random models is found to be associated with strong correlations in the population.

---

The principle of maximum entropy was introduced in 1957 by Jaynes [1, 2] to formulate the foundations of statistical mechanics as an inference problem. Its interest has been recently rekindled by its application to a variety of data-rich fields, starting with the correlated activity of populations of retinal neurons [3, 4]. The method has since been used to study correlations in other neural data, such as cortical networks [5, 6] and functional magnetic resonance imaging [7], as well as in other biological and non-biological contexts, including multiple sequence alignments of proteins [8–10] and nucleic acids [11, 12], the collective motion of bird flocks [13], the spelling rules of words [14], and the statistics of decisions by the United States supreme court [15]. In many cases, the close link between maximum entropy and statistical mechanics has led to new insights into the thermodynamics of the system in terms of phase transitions [16–18], or multi-valley energy landscape [19, 20]. In other cases, the method has allowed for predictions of crucial practical relevance, such as residue contacts in proteins [21], or deleterious mutations in HIV [22].

Although the motivations of maximum entropy seem intuitive and can be formalized rigorously [23], the perceived arbitrariness of its assumptions has led to question its validity [24, 25]. The starting point is to consider models that match empiral observations on a few key statistics of the data. Maximum entropy's crucial—and arguably debatable—assumption is to pick, out of the many models that satisfy that constraint, the one with the largest Gibbs entropy. This choice seems natural, since it ensures that the model is as random as possible. However, it is not clear why it should describe the data better than other models satisfying the same constraints. To address this question directly on empirical data, we reanalyse the original neural data from [3], which contributed to the recent surge of interest in maximum entropy. We compare the accuracy of maximum entropy distributions to ensembles of distributions that satisfy the same constraints, using the approach developped in [26, 27].

The collective state of a population of *N* variables is described by *σ* = (*σ_1_*,…, *σ_N_*). In general, *σ_i_* may denote any degree of freedom, such as the identity of an aminoacid in a protein, the orientation of a bird in a flock, etc. To fix idea, in this paper *σ_i_* will be a binary variable describing the spiking activity of neuron *i*: *σ_i_* = 1 if neuron *i* spikes within a given time window, and 0 otherwise. The joint distribution of the collective activity *σ*, denoted by *P*(*σ*), lives in a 2^*N*^ − 1 dimensional space, represented schematically in Fig. 1. Because that space is huge for even moderately large populations, it is often impossible to sample the true distribution, 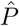, reliably from the data. Simplifying assumptions are needed.

**FIG. 1:**
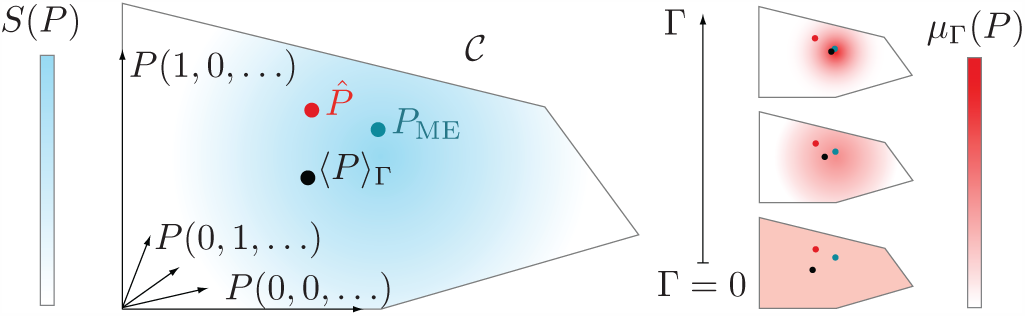
Random models. The space *C* of models *P* is a simplex of 2^*N*^— *M* − 1 dimensions, defined by the intersection of the hyperplanes satisfying the constraints that the mean observables under the model, ∑_*σ*_ *P*(*σ*)𝒪_*a*_(*σ*), *a* = 1,…,*M*, equal the empirical means, 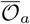, and by a normalization and positivity constraint. The true distribution to be approximated, 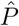 (red dot), is not accessible in general. An entropy-dependent measure *μ*_Γ_ (Eq. 1) is defined on *C* (red map). At Γ = 0 (random ensemble), the measure is uniform over that space. As Γ is increased, the measure concentrates onto the maximum entropy distribution *P*_ME_ (blue dot), and so does its mean *P*_Γ_ = 〈*P*〉_Γ_ (black dot).

To restrict the search of models, one can focus on distributions that agree with the data on the average value of a few observables. Calling these observables 𝒪_*a*_(*σ*), *a* = 1,…,*M*, the condition reads 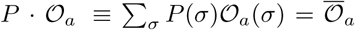, where 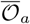 is the empirical mean. The observables must be chosen carefully depending on the problem at hand, and may include local or global order parameters, marginal probabilities, correlation functions, etc. Let us denote by *C* the subspace of models *P* that satisfy those constraints, as well as the conditions *P*(*σ*) ≥ 0 and ∑_*σ*_ *P*(*σ*) = 1. *C* is convex because of the linear nature of the contraints.

A probability law on *C* may be defined which weighs models *P* ∈ *C* according to their Gibbs entropy, *S*(*P*) = -∑*_σ_ P*(*σ*) log *P*(*σ*), through the following measure [26]:

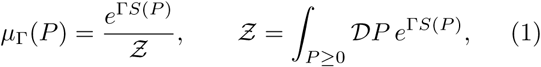

with

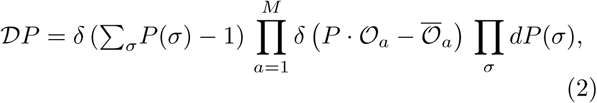

where *δ*(·) is Dirac's delta function. The parameter Γ is conjugate to the entropy, and sets its average value: 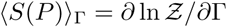, where we use the brackets 〈·〉_Γ_ for averages over the measure *μ*_Γ_. Γ plays the same role with respect to the entropy as the inverse temperature with respect to the energy in standard statistical mechanics. When Γ = 0, all distributions satisfying the constraints have the same probability. We call this the unbiased ensemble. As Γ→ ∞, the measure becomes increasingly peaked onto a single distribution, *P*_ME_, of maximum entropy (or, in the previous analogy, the ground state reached at zero temperature). This distribution defines the classical maximum entropy model, and takes the form [28]:

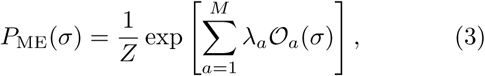

where λ_*a*_ are Lagrange multipliers enforcing the constraints on the mean observables, and *Z* is a normalization constant. We define the average distribution as *P*_Γ_(*σ*) ≡ 〈*(P*(*σ*)〉_Γ_, which belongs to *C* by convexity. *P*_Γ_ only coincides with *P*_ME_ in the limit Γ → ∞. At the other extreme, *P*_0_ is the unbiased, center-of mass distribution that satisfies all the constraints.

We follow the approach of the random ensemble defined by (1) to describe the joint spiking activity of retinal ganglion cells reported in [3]. There, the spiking activities of 40 ganglion cells from the salamander retina were recorded by multielectrode arrays for about an hour, and segmented into ≈ 1.5 · 10^5^ binary spike words *σ* of 20ms. The collective behavior of small networks (up to 10 neurons) was shown to be well described by maximum entropy distributions constrained by spike rates and pairwise correlations (and later to much larger populations [20]). This choice of constraints corresponds to the observables 𝒪_*a*_ = *σ_i_* for all neuron *i*, and 𝒪_*a*_ = *σ_i_σ_j_* for all pairs *i*,*j*, for which the maximum entropy distribution (3) takes the form of a disordered Ising model,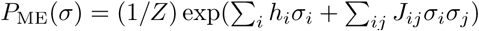.

It is instructive first to consider the unbiased measure *μ*_0_ over very small networks, for which everything can be calculated analytically. The simplest case of two neurons constrained by just their firing rate is illustrated by Fig. 2a. The maximum entropy distribution factorizes over the two neurons, which are thus independent [29]: *P*( *σ*) = *p*_1_( *σ*_1_)*p*_2_( *σ*_2_). Byconstrast, random models drawn from *μ*_0_ are biased towards a positive correlation 〈 *σ*_1_*σ*_2_〉—〈 *σ*_1_〉〈*σ*_2_〉> 0 when both firing rates 〈*σ*_1_〉,〈*σ*_2_〉 are on the same side of 0.5 (in the retinal data 〈*σ*_1_〉 ~ 0.02). A similar bias in the triplet correlation is also found when considering 3 neurons constrained by uniform firing rates and pairwise correlations (Fig. 2b). When pairwise correlations are weak, as is the case in the retina [3], random models predict on average a higher 3-point connected correlation than maximum entropy, although the bias is reversed for large correlations.

**FIG. 2:**
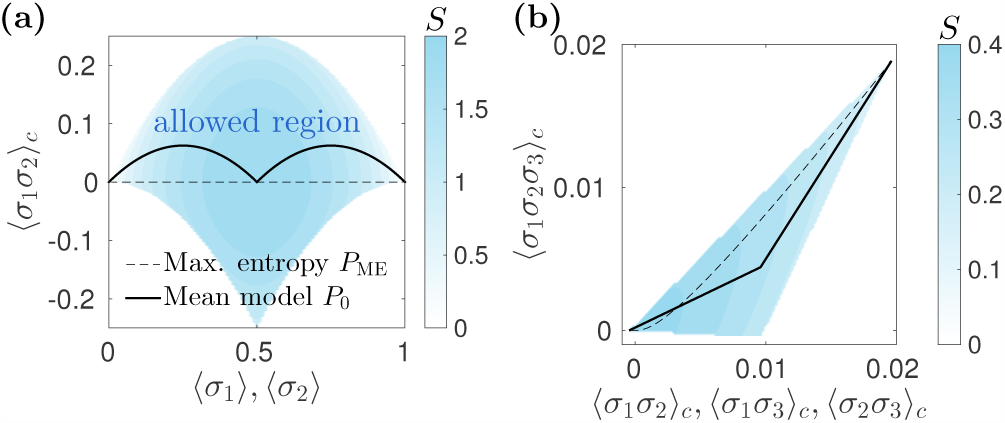
Small networks. Illustration of the random ensemble on 2 and 3 neurons. (a) Pairwise correlation 〈σ_1_σ_2_〉_c_ = 〈σ_1_σ_2_〉 − 〈σ_1_〉〈σ_2_〉 predicted by maximum entropy and random models constrained by the mean spiking rates of two neurons, 〈σ_1_〉 = 〈σ_2_〉, as a function of that rate. The mean unbiased model *P*_0_ is the center of mass between the lower and upper allowed limits of the correlation, which delimit the shaded area. (b) Triplet connected correlation, 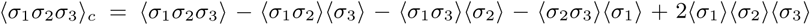 as a function of the pairwise correlation between 3 neurons firing with probability 〈σ_1_〉 = 〈σ_2_〉 = 〈σ_2_〉 = 0.02 (mean empirical value). Pairwise correlation in the retinal data range from −10^−3^ to 0.03 with a median of 2 · 10^−4^. Key is as in (a).

Thanks to its exponential form (3), the maximum entropy distribution can be inferred with relative ease for systems of size *N* ≤ 20, yet requiring to calculate sums of 2^*N*^ terms [3]. Sampling from *μ*_Γ_ or calculating *P*_Γ_, on the other hand, is a much harder task, involving the exploration of *C* of dimension 2^*N*^ − *N*(*N*+1)/2 − 1. To apply the random ensemble to populations of neurons, we sampled from *μ*_Γ_ using the Metropolis-Hastings algorithm, for various subgroups of neurons of different sizes. At each step, starting from a distribution *P* in *C*, one picks a random direction *V* in the Fourier basis of the hyperplane orthogonal to all observables 𝒪_*a*_ [27]. The new distribution is taken to be *P′* = *P* + *αV*, where *α* is drawn uniformly irval (*α*_min_,*α*_max_) defined by the lower and upper limits so that *P*′(*σ*) ≥ 0 for all *σ*. *P*′ is accepted with probability min(1,*e*^Γ[*S*(*P*′)—*S*(*p*)]^). The process is repeated until equilibration is reached. High space dimension limits us to relatively small group sizes, *N*≤ 8. Fortunately for these sizes the true distribution 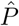 may be accurately estimated from the data, and directly compared to models.

The accuracy of a given model is assessed by the Kullback-Leibler (KL) divergence between the model distribution *P* and the true one 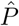, 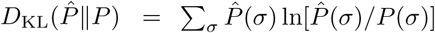. Fig. 3 shows, in the form of violin plots, the distribution of KL divergence (normalized relative to maximum entropy) when sampling *P* from *μ*_Γ_, for groups of *N* = 7 cells. This distribution is plotted in Fig. 3a for a random groups of 7 cells. Maximum entropy is found to have a clear advantage: its accuracy is matched by only a negligible fraction of models drawn from *μ*_Γ_, and it also does better than their mean *P*_Γ_ (red line). The advantage of maximum entropy over the unbiased ensemble generalizes to 20 random groups of 7 cells (Fig. 3b), as well as the groups comprising the most correlated (green) and most active (yellow) cells. Interestingly, in all cases the mean distribution *P*_Γ_ is more accurate than the typical distribution *P* sampled from *μ*_Γ_, a consequence of Jensen's inequality which implies 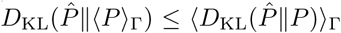. In general, 0 < Γ<∞ to interpolates between the unbiased ensemble and the maximum entropy distribution. For these reasons, in the following the maximum entropy model *P*_ME_ will only be compared to the mean distribution of the unbiased ensemble, *P*_0_.

**FIG. 3:**
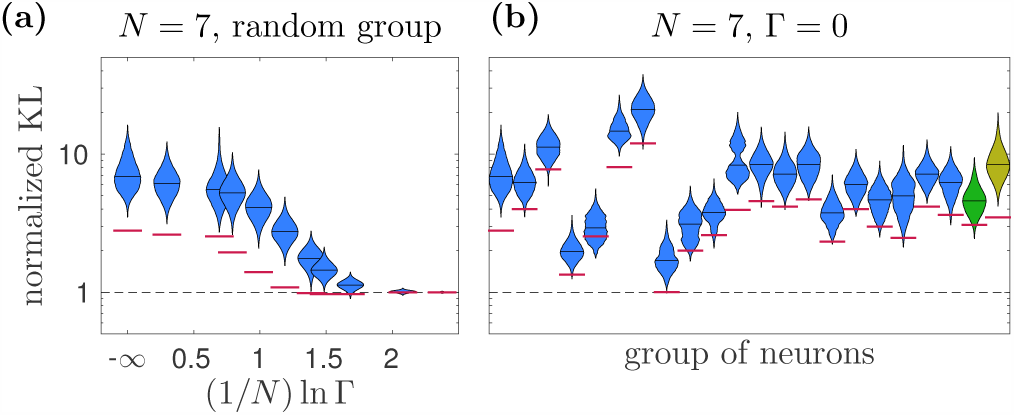
Random versus maximum entropy models. (a)The normalized Kullback-Leibler divergence relative to maximum entropy, 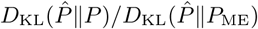, is represented as a function of the entropy-conjugated variable Γ for (a) a random group of *N* = 7 neurons (out of 40). Values above unity (dashed line) mean that maximum entropy outperforms the random model. The violin plots show the distributions over random models drawn from *μ*_Γ_, while the red lines show the value for the average model, 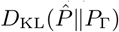. (b) Normalized KL divergence at Γ = 0 for 20 random subsets of 7 neurons (blue), as well as the group of most correlated neurons (as measured by Pearson's correlation coefficient, green), and the set of neurons with the highest spike rate (yellow).

We now investigate the dependence on the population size. Fig. 4 shows the average normalized KL divergence of the mean model *P*_Γ_ for random cell groups of varying sizes, as a function of (1/*N*)ln Γ (the scaling of Γ is assumed to be exponential in *N*, as suggested by calculations with random observables [26]). The general trend noted before for *N* = 7 generalizes to all sizes: the larger the entropy bias Γ, the better the model (Fig. 4a). However, this average behaviour masks large heterogeneities across different choices of cell groups, especially for small groups, of which a sizeable fraction is better described by the mean distribution *P*_0_ than by *P*_ME_. Evaluating this fraction from hundreds of random groups for each *N*, we find that maximum entropy is more likely to outperform the random ensemble in larger groups (Fig. 4b), and even does so in all of the 200 tested groups of size *N* = 8.

**FIG. 4:**
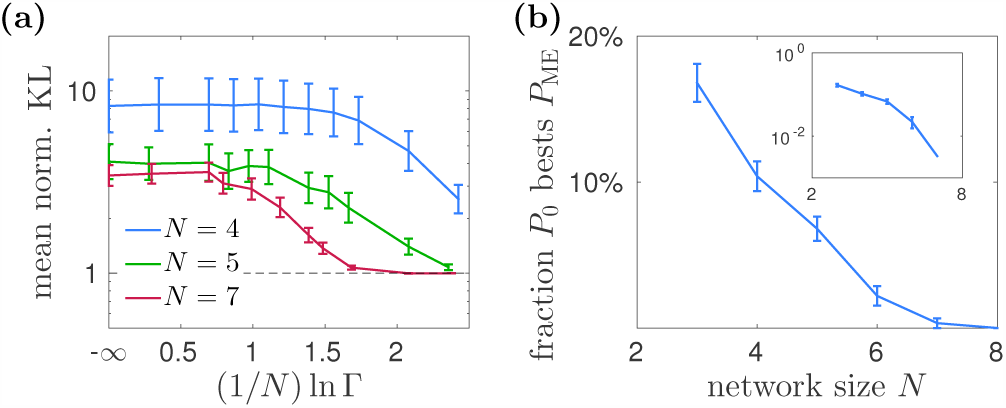
Dependence on populations size. (a) The normalized divergence of the average model, 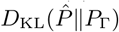, is averaged over 20 random subsets, and plotted as a function of (1/*N*)ln Γ. Errors bars show standard error on the mean. (b) Fraction of random groups (out of hundreds) of *N* neurons that are better described by the mean unbiased distribution *P*_0_ than by the maximum entropy model *P*_ME_.

What sets apart groups of cells that are better described by *P*_0_ than by *P*_ME_? Since both share the same 1- and 2-point correlations by construction, we examine their predictions for 3-point correlations in triplets of cells (*N* = 3). Fig. 5a shows that random models typically fail because they overestimate small 3-point correlations. By contrast, maximum entropy is more likely to be outperformed by random models when the triplet correlation is large, in which case maximum entropy overestimates it. Both these findings are in agreement with the results of Fig. 2b. This observation can be generalized to larger groups of neurons (*N* > 3) by considering the total amount of correlations in the network, quantified by the loss of entropy due to correlations, or multi-information [29], 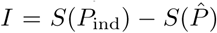, where 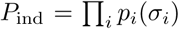 is the model distribution of independent neurons. Groups that are better described by *P*_0_ than by *P*_ME_ are found to have a higher multi-information on average (Fig. 5c).

**FIG. 5:**
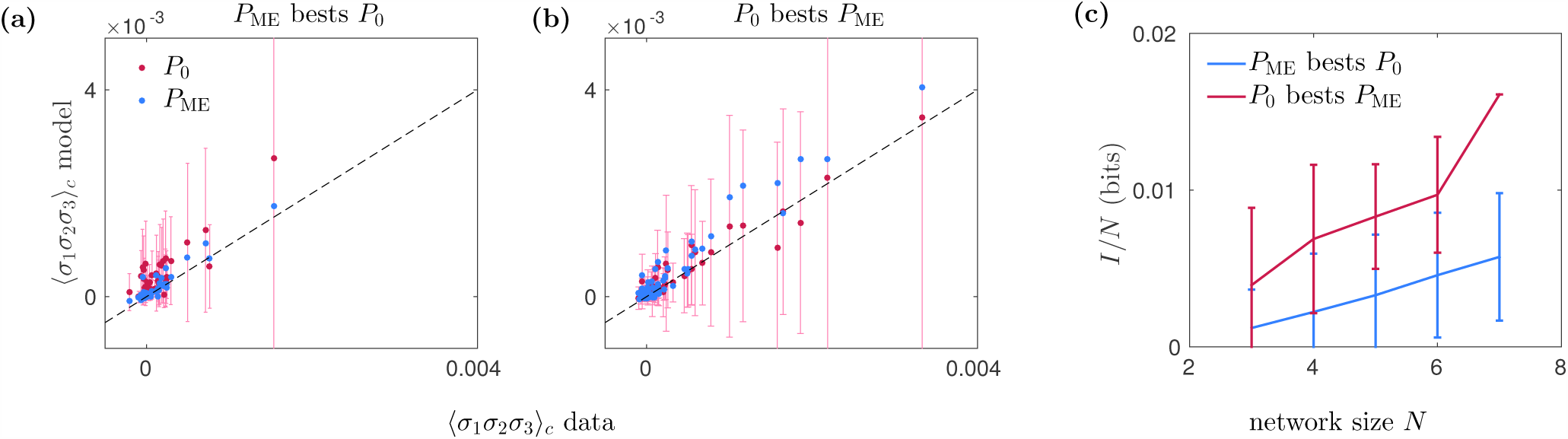
Correlations and maximum likelihood performance. Three-point connected correlation 〈*σ*_1_*σ*_2_*σ*_3_〉_*c*_ for (a) 100 random triplets whose joint activity is best described by maximum entropy and (b) 100 random triplets whose joint activity is best described by the mean unbiased model, when constraining the values of the pairwise correlations. The error bar shows, for each triplet, the allowed range of values for the 3-point correlation. (c) The multi-information, which measures the overall amount of correlation in the collective acitivity, is plotted as a function of system size, for groups of neurons that are best described by the maximum entropy model *P*_ME_ (blue) or by the mean unbiased model *P*_0_ (red). Error bars show standard deviation across groups of cells (the red point at *N* = 7 has no error bar because only one group of that size was better described by *P*_0_).

Since maximum entropy was proposed as a method for building statistical models from high-dimensional data, its accuracy, relevance, and epistemological validity have been questioned. In this study we have shown that the maximum entropy model describes the spiking activity of populations in the retina better than the mean model satisfying the same constraints, which itself performs better than the vast majority of random models under these constraints. This better performance of maximum entropy gets more marked as the population size *N* grows, and is essentially always true for *N* ≥ 8. The analysis of 3-point and higher-order correlations suggests that the rare instances where the mean model outperforms maximum entropy is when correlations are relatively large. In that case, maximum entropy predicts high triplet correlations within the allowed range compared to the mean unbiased model (Fig. 2b), and may thus overestimate their true value, consistent with previous observations in large populations [20]. In that case, models that take a “middle-of-the-road” value of the correlations may be preferred to maximum entropy.

By providing a test on empirical data, our results complement previous work aimed at explaining or refuting the efficiency of maximum entropy based on theoretical arguments and simulated datasets. Calculations on synthetic learning problems have suggested that maximum entropy is no more accurate than random [26], unless the chosen observables are smooth as a function of configuration space [27]. However, in these studies the choice of observables and true distributions were taken to be completely random, and it is not clear how applicable they are to real distributions and to pairwise constraints.

Other simulation studies have more specifically addressed the role of pairwise interactions. It was suggested that pairwise maximum entropy models should fail for large populations [30]. On the other hand, strongly interacting systems with interactions of arbitrary order have been numerically shown to be well described by pairwise interactions, with an analogy to Hopfield networks [31]. The principle of maximum entropy has also been advocated by contrast to non-additive (or Rényi) entropies, but on purely theoretical grounds [32]. Our results do not preclude that other objective functions than entropy may help better describe empirical data. They suggest, however, that it is better to pick the most random model than to pick a model at random.

We thank Michael Berry for sharing the data from [3], and Olivier Marre for his comments on the manuscript.

